# Structure-Guided Biochemical Design of DNA Tweezers As A Dual Target of the Primary Glioblastoma Biomarkers S100A4 and Midkine

**DOI:** 10.64898/2026.04.26.720909

**Authors:** Hayley Foo, Gaurav Sharma

## Abstract

Glioblastoma multiforme (GBM) is among the most aggressive malignant brain tumors originating from glial cells and characterized by severe infiltration into surrounding brain tissue, rendering early detection difficult with current diagnostic imaging methods. S100A4 has been identified as a biomarker protein associated with glioblastoma invasiveness due to its role in cell motility and tumor metastasis. Similarly, midkine (MDK) poses an optimal biomedical target for identifying GBM invasive phenotypes because of its connection to the tumor microenvironment and infiltrative proliferation. Both proteins notably possess a positive charge that interacts electrostatically with the negatively charged phosphate backbone of DNA. It has been established that early molecular detection remains a critical unmet need. This study investigates a promising strategy for GBM diagnosis based on how S100A4 and MDK can selectively bind with DNA tweezer nanostructures. Computationally predicting eight distinct nucleotide sequences yielded three-stranded, hinge-scaffolded tweezer conformations for each candidate. The target protein and DNA structures, derived from AlphaFold, were paired together by molecular docking simulations conducted with HDOCK. Docking analyses evaluated binding affinity, structural complementarity, and conformational stability of the complexes formed. Among the evaluated candidates, DT3_8 computationally established the most biochemically robust interaction with both biomarker proteins. Selectivity is especially important because many S100 proteins share similar electrostatic profiles, yet DT3_8 indicates stronger selectivity for S100A4 and MDK over other S100 family proteins. These findings establish a biomechanical basis for the development of nanoscale DNA biosensors, which suggests the potential for detecting invasive GBM phenotypes, preceding radiographic manifestation and pending experimental validation.

## 1. Introduction

Glioblastoma multiforme (GBM) is the most aggressive and lethal primary malignant brain tumor in adults. Characterized by rapid proliferation and extensive infiltration into surrounding healthy brain tissue ***[1]***, GBM is classified as a grade IV astrocytoma with very poor prognosis. The current standard of care, including surgical resection, radiation therapy, and chemotherapy, remain largely ineffective as the median survival ranges between 15-18 months and 5-year survival is approximately lower than 10% ***[2]***.

Current imaging-based approaches, including MRI and PET, provide anatomical information but fail to detect early molecular tumorigenic changes ***[3,4]***. GBM exhibits structural heterogeneity and diffuse infiltration in regions inherently resistant to contrast-enhancement, making complete surgical resection nearly impossible ***[3]***. Moreover, imaging modalities, namely, MRI, often struggle to distinguish true tumor progression from pseudoprogression, defined as the phenomenon when treatment combinations such as temozolomide chemotherapy and radiation indirectly and temporarily initiate tumor worsening and radiation necrosis ***[4]***. Thus, a critical diagnostic gap exists between current structural imaging techniques and the molecular changes that drive early GBM progression.

Among molecular drivers of GBM invasion, the calcium-binding protein S100A4 (metastasin-1) has emerged as a key biomarker ***[5]***. S100A4 regulates cytoskeletal remodeling, epithelial-mesenchymal transition-like signaling, and metastatic pathways ***[6]***. Elevated S100A4 expression also correlates with enhanced motility, increased metastatic potential, and poor clinical prognosis ***[6]***. Structurally, S100A4 is enriched in lysine and arginine residues, producing a strongly positive electrostatic surface ***[7]***. This charge distribution makes S100A4 an attractive biochemical target for negatively charged nucleic-acid nanostructures.

As aligned with the hypothesized dual function of DNA tweezers as a nano-scale detection device for glioblastoma, midkine (MDK) was also identified as a biomarker protein associated with GBM’s infiltrative nature and a driver of proliferation ***[8]***, as evidenced by its extracellular location and function as a secreted protein from an initial UniProt identification. One of its most indicative features is that a surplus of midkine is linked to the mutated EGFRvIII protein, a key genotype that precedes GBM invasion and drives signaling pathways connected to GBM tumor progression ***[9]***. For one, MDK is also a mechanistic catalyst for driving macrophage proliferation and expanding the tumor microenvironment ***[10]*** and is inherently positively charged at established core binding residues.

DNA nanotechnology offers a programmable and versatile platform for molecular sensing and targeted therapeutic delivery. DNA tweezers are mechanically responsive nanostructures assembled from synthetic oligonucleotides that transition between open and closed states upon target binding ***[11]***. The negatively charged phosphate backbone of DNA enables electrostatic attraction to positively charged proteins such as S100A4 and MDK. In addition, DNA tweezers, essentially assembled at the nano-scale, can be functionalized with fluorescent reporters or therapeutic payloads, enabling diagnostic and theranostic applications ***[12]***.

Developing a molecular-scale sensor that can dually recognize S100A4 and MDK, all in all, aims to address limitations of current structural imaging approaches. Because S100A4 and midkine overexpression reflects invasive tumor behavior before gross anatomical changes appear and can be tracked radiologically, a nanosensor capable of detecting this protein may provide earlier, biologically relevant diagnostic signals, if validated experimentally. By designing DNA tweezers that selectively recognize these biomarker proteins, this work explores a biochemical detection strategy that operates at the molecular level, potentially laying the groundwork for low-cost, scalable, and non-invasive diagnostic tools for aggressive cancers similar to GBM.

## 2. Methods

The project followed a structured, multi-phase computational outline designed specifically to move from raw, sequenced data to an *in silico*-validated protein-DNA nanodevice complex. Each phase built upon the previous one, combining systematic processes such as sequence retrieval, structure prediction, molecular docking, interaction analysis, and molecular dynamics simulations.

### 2.1 Phase I: Protein Modeling

The amino acid sequence of S100A4 was retrieved from the UniProt database with an ID of P26447 ***[13]***: [MACPLEKALDVMVSTFHKYSGKEGDKFKLNKSELKELLTRELPSFLGKRTDEAAFQKLMSNLDSNRDNEVDFQEYCVFLSCIAMMCNEFFEGFPDKQPRKK] and used as one primary input for all further exploration. The MDK amino acid sequence was also identified similarly: [MQHRGFLLLTLLALLALTSAVAKKKDKVKKGGPGSECAEWAWGPCTPSSKDCGVGFREGTCGAQTQRIRCRVPCNWKKEFGADCKYKFENWGACDGGTGTKVRQGTLKKARYNAQCQETIRVTKPCTKTKAKAKAKKGKGKD]. AlphaFold was employed to simulate the three-dimensional structures of S100A4 and MDK using the amino acid sequence inputs and would also offer integral information about internally relevant attention mechanisms quantified by pLDDT confidence scores to identify the reliability and surface accessibility of specific protein regions ***[14]***. The center regions of both S100A4 and MDK, shown in blue, promise the most stable bonding interaction with the DNA tweezer candidates (*Figure 1A, 1B)*.

**Figure 1.**
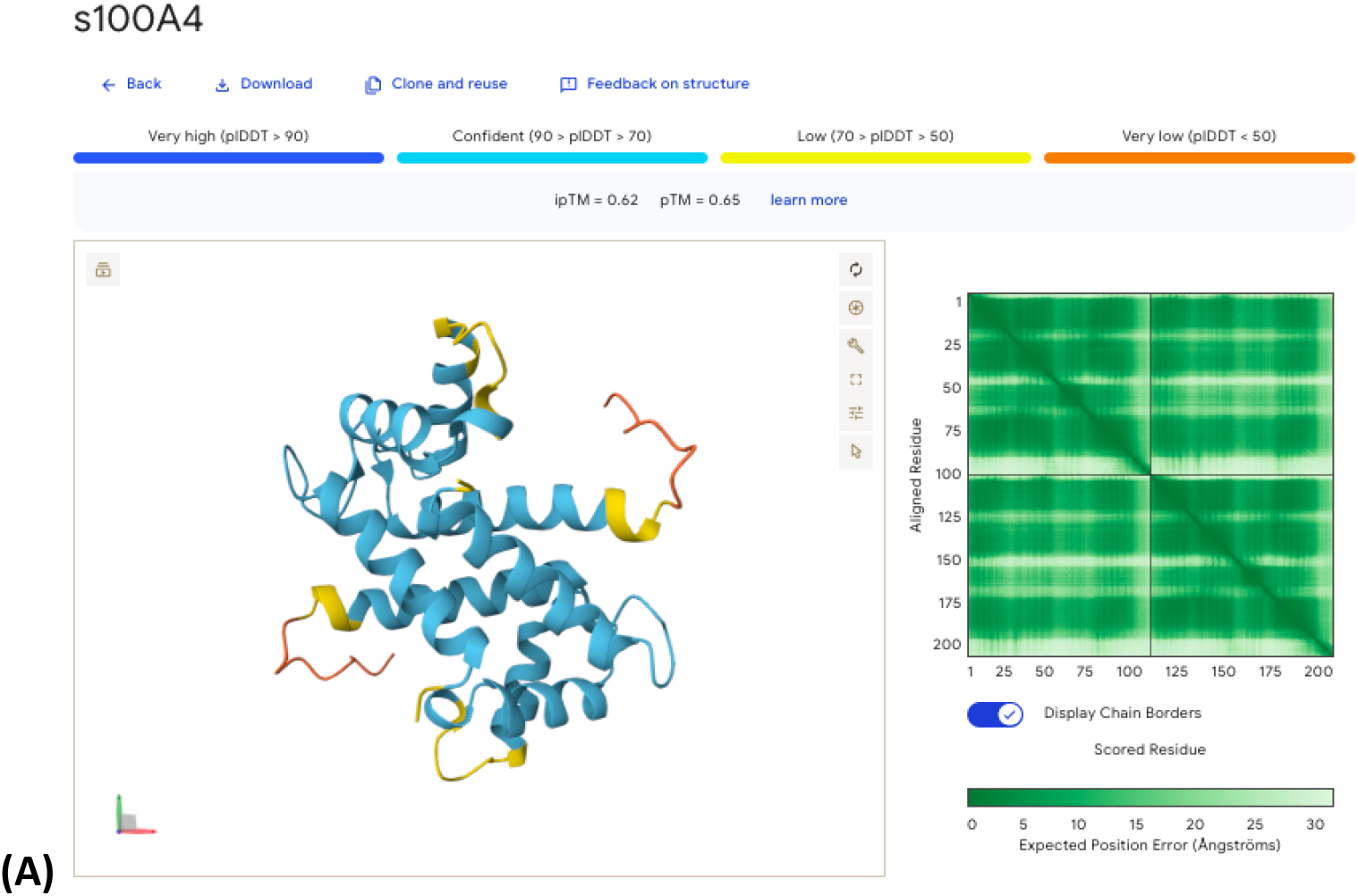
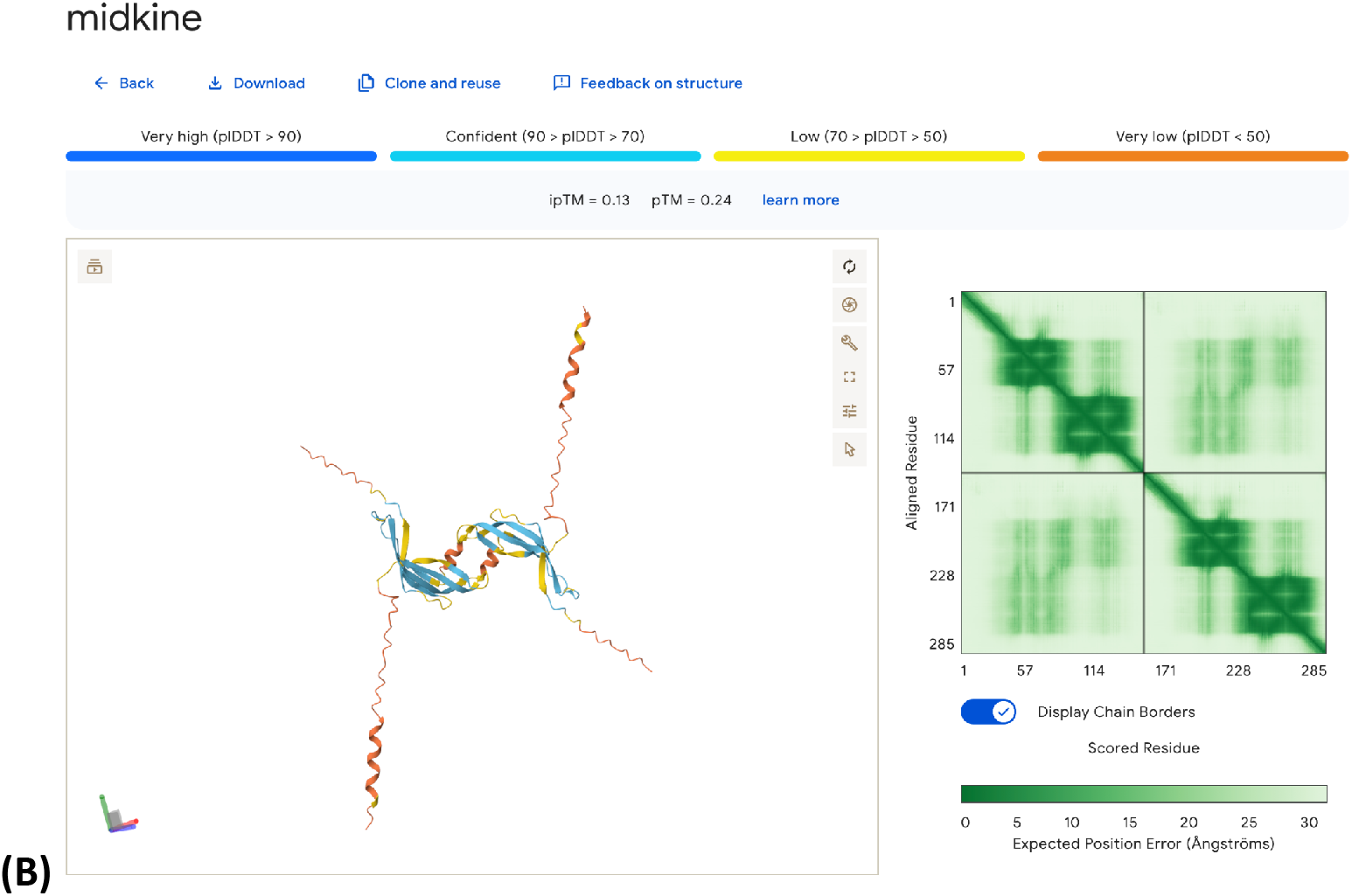
Snapshot of AlphaFold output when software was run to visualize the S100A4 **(A)** and MDK **(B)** structure and their respective key residues.

Regions with lower confidence were visually inspected and simulated for surface structures and shape in ChimeraX ***[15]*** to confirm specific binding pockets identified by P2Rank ***[16]*** at a later time to align with blind docking processes, where alternative softwares’ determined optimal binding sites have no effect on researchers’ findings or predictions ***[17]***. The final S100A4 and MDK structures were then exported in CIF format, converted to PDB on ChimeraX, and designated as the receptor for all subsequent docking simulations and phases.

### 2.2 Phase II: DNA Tweezer Design

A baseline DNA tweezer design consisting of three strands (*Figure 2)* was selected as the starting template for which open- and closed-state configurations could be manually designed, deemed DT0.

**Figure 2.**
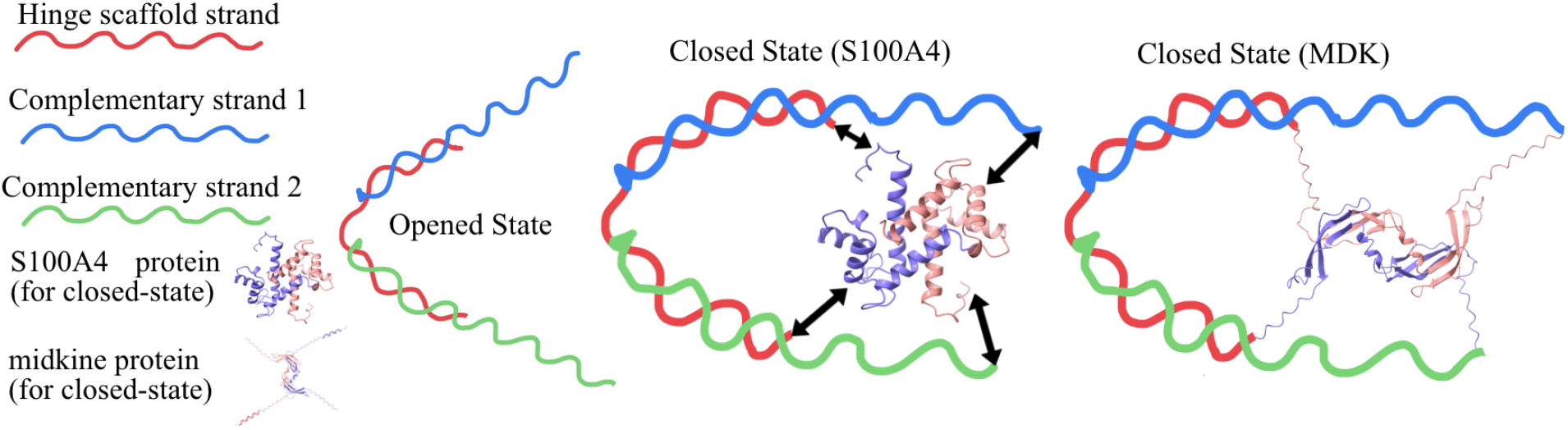
Schematic representation of DNA tweezer folding, with functional open-closed mechanisms. Binding of S100A4 and MDK uniquely induce tweezer closure.

To improve mechanical rigidity and folding stability, the arms of the tweezer were elongated by 1-2 codons (triplets of nucleotides), generating a set of eight distinct candidate sequences. Each candidate sequence was then input into AlphaFold to predict its three-dimensional conformation. The resulting models were examined for proper folding, symmetry between arms, and any extraneous presence of severe distortions or unintended secondary structures. Candidates that displayed well-formed, stable open conformations with clearly defined arms were retained for docking. These DNA structures were also exported in PDB format for use as ligands in the docking phase.

### 2.3 Phase III: Molecular Docking & Interaction Profiling

In the docking phase, each DNA tweezer candidate was paired with the S100A4 and MDK structures in independent binding events simulated in HDOCK ***[18]***. For each protein-DNA pair, HDOCK generated a set of ten predicted docking models representing different possible binding orientations and interfaces. To rank every model, HDOCK reported a docking score that reflected the software’s confidence in binding predictions, as well as a ligand RMSD and an overall confidence score. These metrics were collected and compared across all eight DNA tweezer complexes. Complexes with lower (more negative) docking score values were interpreted as having stronger predicted binding affinity ***[19]***. Consistently low ligand RMSD values across the top models for a given tweezer indicated a stable and reproducible binding capacity. Based on these criteria, the most promising candidate would be selected for future analysis beyond the scope of this particular study.

To understand the biochemical basis of the docking results, the top-ranked complexes from HDOCK were analyzed using PLIP ***[20]***. For each selected protein-DNA complex, PLIP scanned the interface and identified specific non-covalent, crucial biochemical interactions, including hydrogen bonds, hydrophobic contacts, and salt bridges. Hydrogen bonds were evaluated based on donor-acceptor distances and angles, hydrophobic contacts were identified between non-polar residues and nucleobases or sugar-phosphate regions, and salt bridges were detected between positively charged amino acid side chains (such as lysine and arginine) and negatively charged phosphate groups on the DNA backbone. The number, type, and spatial distribution of these interactions were compared across candidates. Complexes with a higher quantity of viable hydrogen bonds and salt bridges, particularly involving S100A4’s and MDK’s basic residues, respectively, were interpreted as having stronger and more specific electrostatic and structural complementarity based on the extensive hydrophobic interactions between the two subjects within the protein-DT complex candidates.

### 2.4 Phase IV: Molecular Dynamics Simulations

To evaluate whether the most promising complexes would remain stable under conditions approximating the human body, molecular dynamics simulations were performed using GROMACS ***[21]***. The selected protein-DNA complex would be placed in a 3D simulated environment with water molecules to mimic an aqueous physiological environment unique to the human body ***[22]***. Appropriate ion concentrations, namely, sodium and chloride, were added to neutralize the system and approximate physiological and naturally occurring ionic strength ***[23]***. A production run was then carried out over a defined simulation time.

Throughout the simulation, root-mean-square deviation (RMSD) was calculated to track overall structural drift of the complex over time, while root-mean-square fluctuation (RMSF) was used to measure the flexibility of individual residues and nucleotide positions. Stable RMSD values and low RMSF at the binding interface would indicate that the DNA tweezer maintained its structure and binding interactions with S100A4 and with MDK over time in a water-based environment.

## 3. Results

This multi-phase computational methodology generated a comprehensive evaluation of eight select DNA tweezer candidates and their interactions with the S100A4 protein, then sequentially with the midkine protein. Each stage—structure prediction, docking, interaction profiling, and molecular dynamics—contributed distinct evidence that converged on candidate DT3_8 as the most viable nanodevice.

### 3.1 Structural Modeling of S100A4, MDK, and DNA Tweezers

AlphaFold produced a high-confidence structural model of S100A4, with pLDDT values indicating strong reliability across the EF-hand calcium-binding domains and the positively charged C-terminal region. The same result was obtained with AlphaFold’s simulation of MDK’s structural components. These regions are known to mediate protein-ligand interactions and therefore represent plausible docking interfaces for DNA-based nanostructures. The electrostatic surface map generated in ChimeraX confirmed a pronounced cluster of lysine and arginine residues for S100A4 and MDK (*Figure 3)*, reinforcing the hypothesis that electrostatic attraction would be the initial driver of DNA-protein binding but sustained later by biochemical interactions that form upon conformational-change binding.

**Figure 3.**
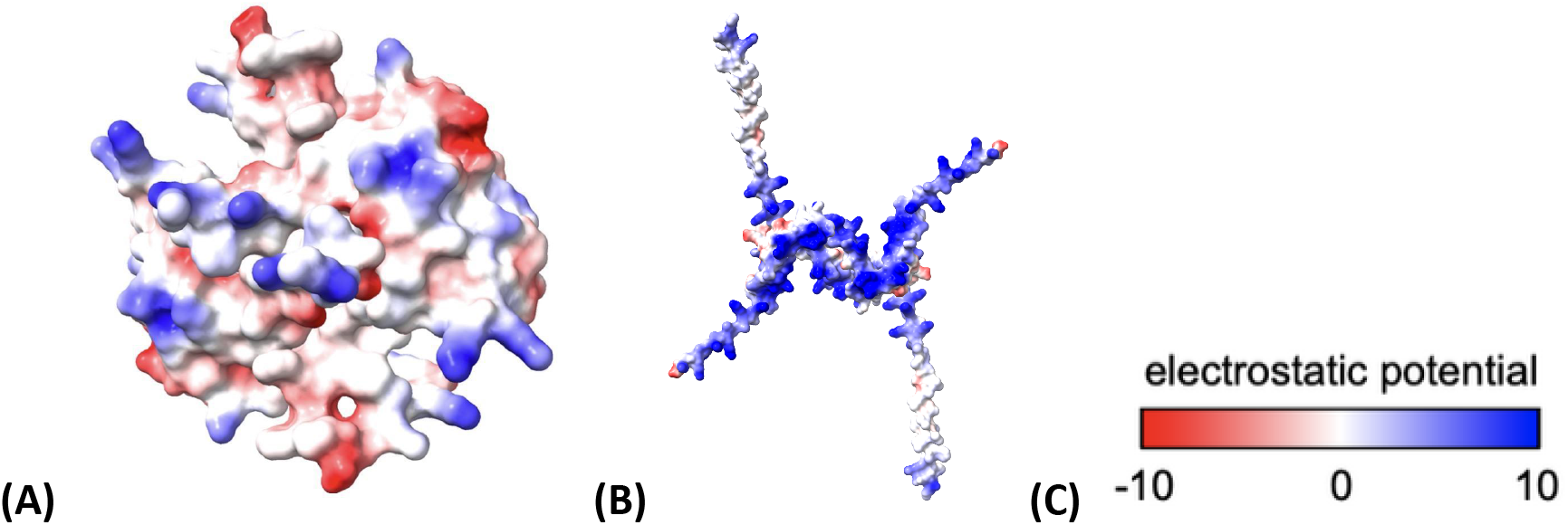
Electrostatic surface potential of S100A4 **(A)** and MDK **(B)**, retrieved from ChimeraX and a legend depicting the difference between red and blue according to UCSF developers **(C)**.

For the DNA tweezers, AlphaFold was used as a platform to run approximately 300 trials over the span of this study to computationally determine viable tweezer structures. Eight were completed and aligned with open-state mechanisms of a DNA tweezer *(Table 1)*. AlphaFold predicted stable conformations for all eight candidates, with visualizations of structural confidence in the tweezers’ inherent open states (*Figure 4)*. However, structural inspection revealed meaningful differences: DT3_8 displayed the most symmetrical arm geometry, minimal strand curvature, and the fewest unintended secondary structures. These features suggested greater mechanical rigidity and more predictable binding behavior, making them strong candidates for subsequent docking analysis, which would be confirmed in the next phase.

**Table 1.**
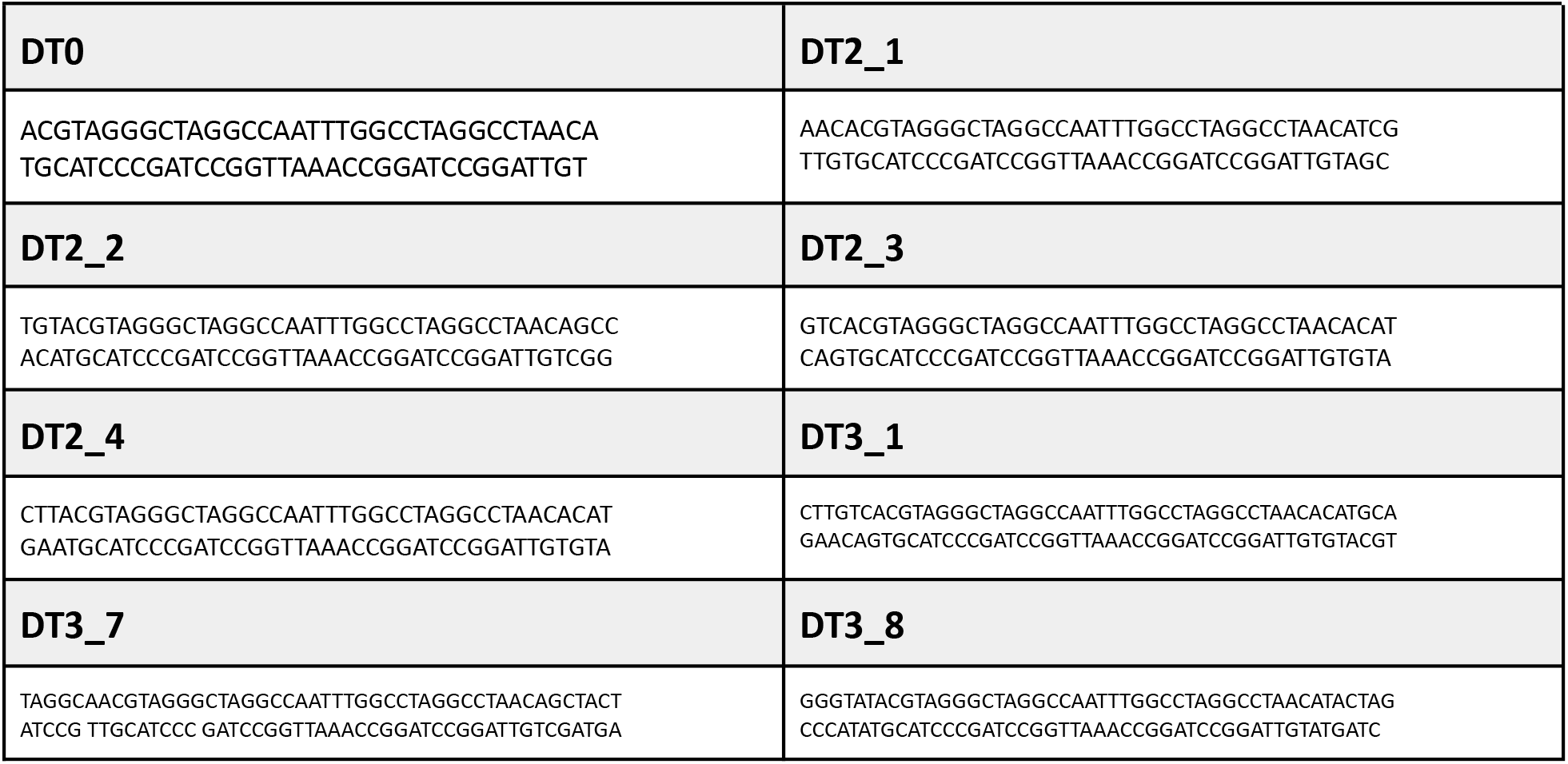
Compilation of eight tweezers’ nucleotide template and complementary sequences, from DT0 to DT3_8.

**Figure 4.**
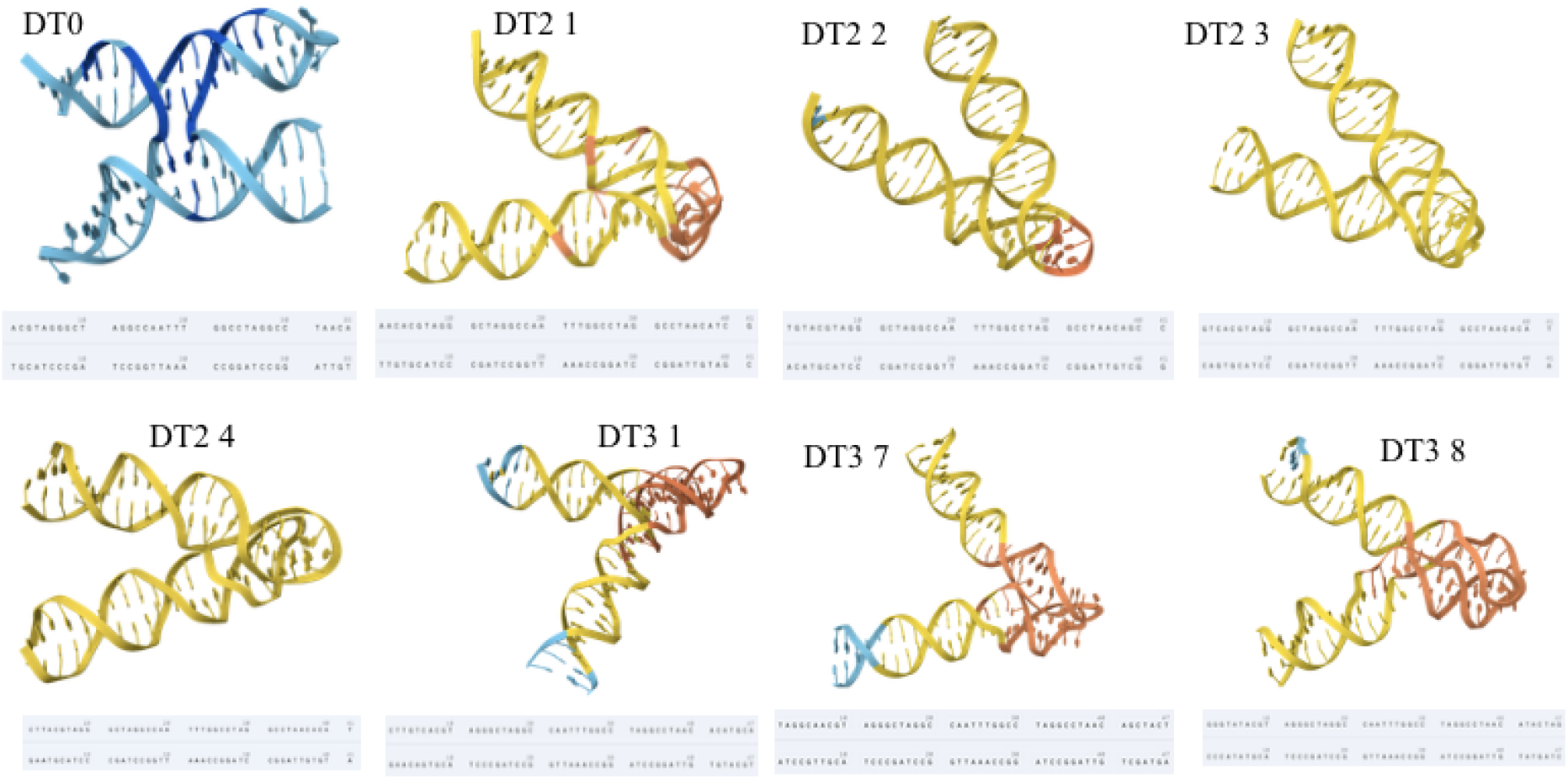
Input of manually designed DNA tweezer nucleotide sequences (Table 1) into AlphaFold to model the 3-D structures of 8 chosen tweezer candidates. Each complete tweezer candidate’s sequence was extended by one codon (DT2 group) and two codons (DT3 group) from the DT0 template model.

### 3.2 Docking Performance and Binding Affinity

Docking simulations using HDOCK revealed clear performance evaluation among the eight DNA tweezers with S100A4 and MDK *(Figure 5, 6 respectively)*. DT3_8 consistently achieved the lowest (most statistically favorable) docking score values across all ten predicted docking positions, indicating the strongest predicted binding affinity to S100A4 and midkine. The docking values for DT3_8 were not only lower than those of other candidates but also showed the highest confidence score (*Table 2)*, a metric given by the equation (*Equation 1*) ***[24]***, suggesting a stable and reproducible binding orientation.

**Table 2.** Docking scores and confidence scores amongst all eight DNA tweezer/S100A4 and DNA tweezer/MDK complexes. A confidence score above 0.7 shows favorable, high likelihood of stable binding **[21]**.

| DNA<br>Tweezers/<br>S100A4<br>Complex | DNA<br>Tweezers/<br>MDK<br>Complex | Best<br>Docking<br>Score<br>(S100A4) | Best<br>Confidence<br>Score<br>(S100A4) | Best<br>Docking<br>Score<br>(MDK) | Best<br>Confidence<br>Score<br>(MDK) |
| --- | --- | --- | --- | --- | --- |
| DT0 (S) | DT0 (M) | -226.19 | 0.8211 | -218.35 | 0.8108 |
| DT2_1 (S) | DT2_1 (M) | -208.05 | 0.7615 | -226.56 | 0.8512 |
| DT2_2 (S) | DT2_2 (M) | -237.34 | 0.8515 | -224.14 | 0.8366 |
| DT2_3 (S) | DT2_3 (M) | -224.49 | 0.816 | -230.66 | 0.8693 |
| DT2_4 (S) | DT2_4 (M) | -226.80 | 0.8229 | -257.22 | 0.8951 |
| DT3_1 (S) | DT3_1 (M) | -219.52 | 0.8007 | -250.71 | 0.8823 |
| DT3_7 (S) | DT3_7 (M) | -218.17 | 0.7963 | -263.12 | 0.9057 |
| DT3_8 (S) | DT3_8 (M) | -256.57 | 0.8939 | -270.84 | 0.9181 |

**Figure 5.**
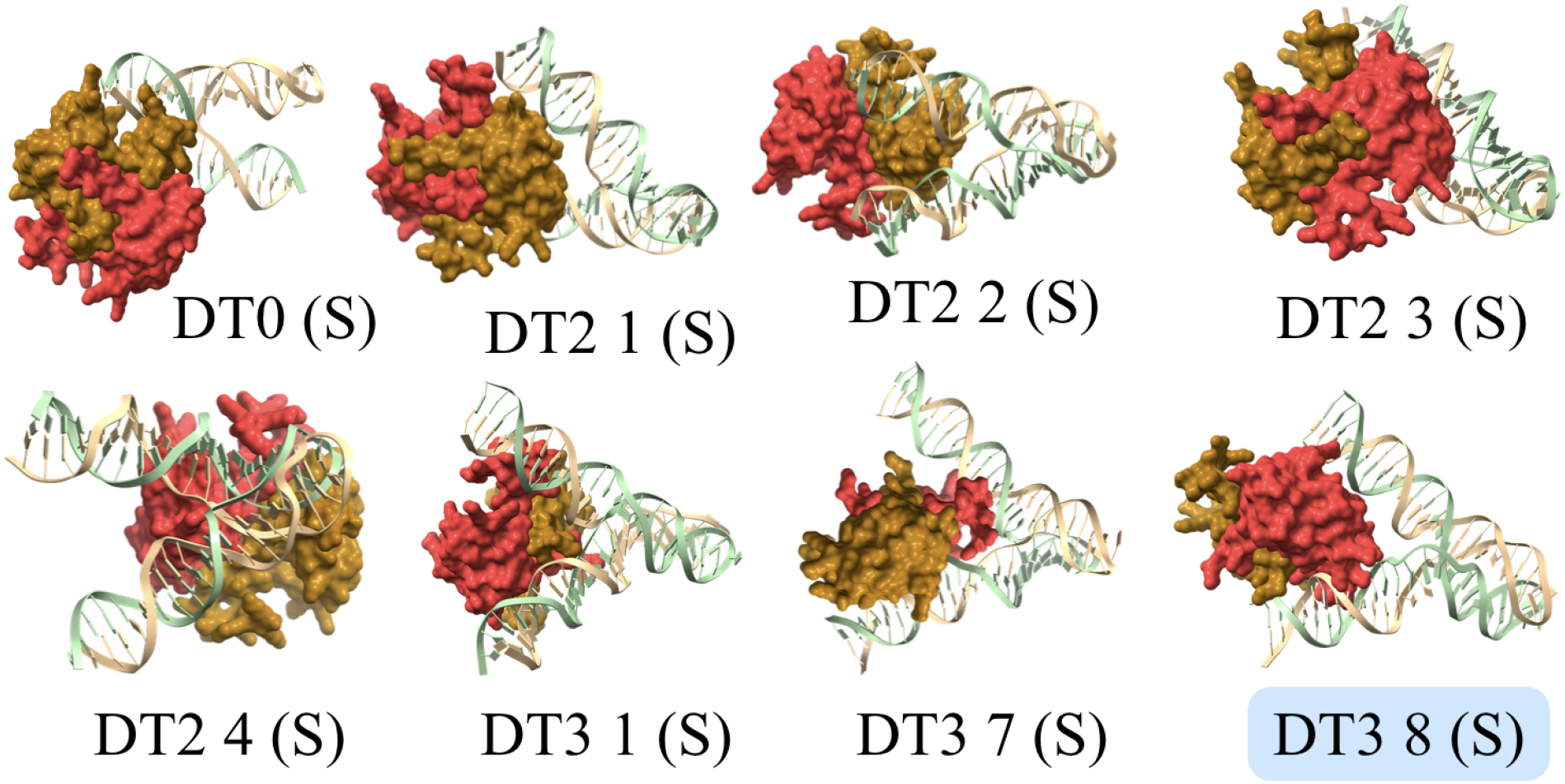
S100A4-tweezer complexes simulated using molecular docking HDOCK software were converted to ChimeraX for viewing. The S100A4 protein is pictured in **red and orange**, DNA tweezer in **pale green and yellow**. Visual and binding statistical analysis determine the DT3_8 candidate to be the best model for future applications and later therapeutic considerations.

**Figure 6.**
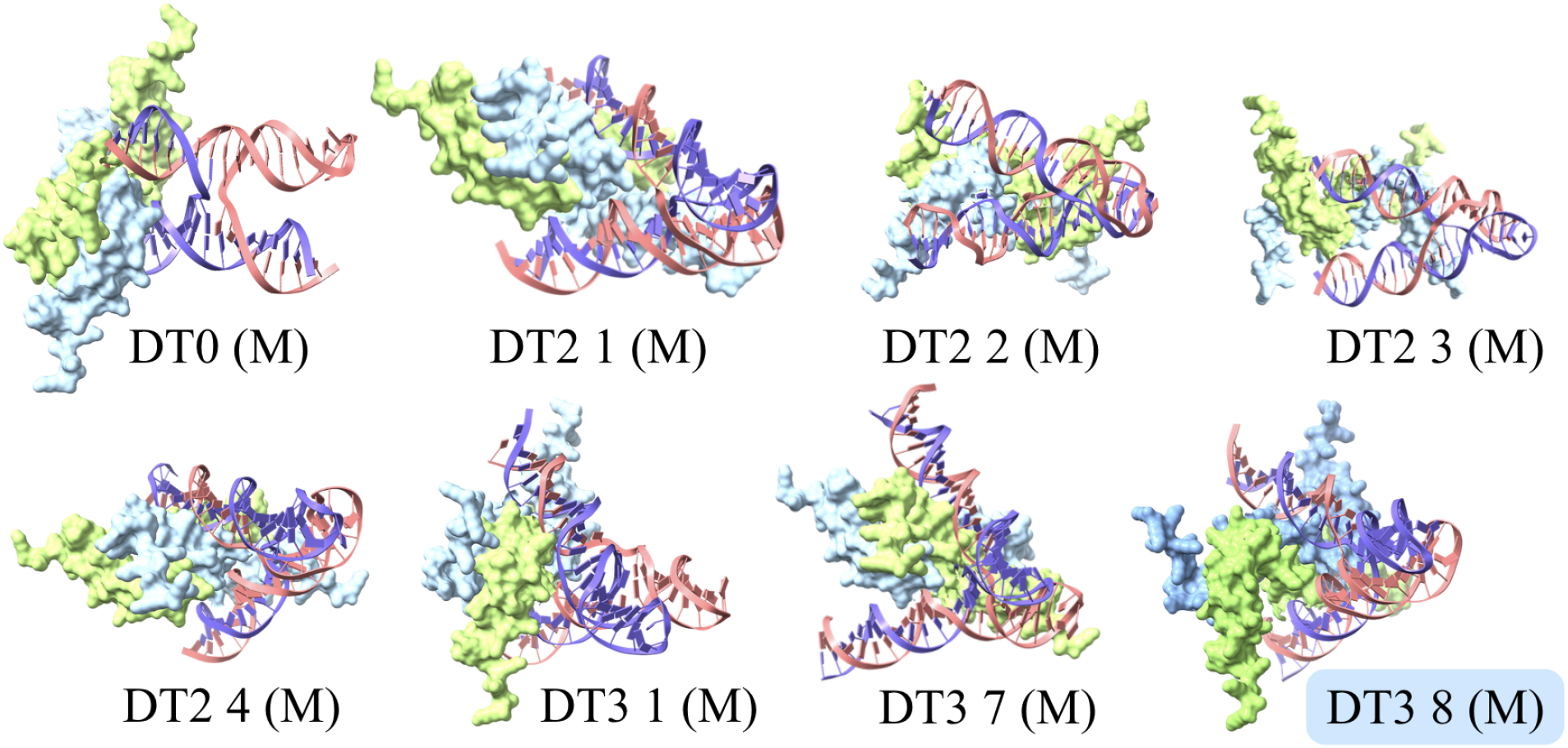
MDK-tweezer complexes simulated using molecular docking HDOCK software were converted to ChimeraX for viewing. The MDK protein is pictured in **green and blue**, DNA tweezer in **pink and purple**. Visual and binding statistical analysis determine the DT3_8 candidate to be the best model for future applications and later therapeutic considerations.

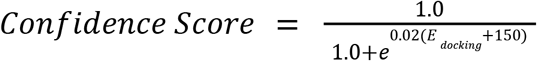

*Equation 1. HDOCK computation formula for confidence score quantities*.

### 3.3 Interaction Profiling and Biochemical Complementarity

Comprehensive interaction profiling using PLIP provided insight into the superior binding performance of the DT3_8 construct relative to other candidate DNA tweezers. Quantitative analysis of both complexes formed with DT3_8 revealed a spatially dense and coordinated network of stabilizing intermolecular binding contacts, indicating that its enhanced docking score was supported by biochemically meaningful interactions.

Findings from PLIP simulations yielded a higher number of viable convergent hydrogen bonding, hydrophobic interactions, and electrostatic salt bridges for DT3_8 *(Figure 7)*, which suggests that it forms a structurally and chemically complementary binding interface with S100A4 and MDK. Enhanced interface specificity and structural persistence as determined are completely consistent with the improved docking scores and binding reproducibility observed for DT3_8, thereby supporting its selection as the lead DNA tweezer candidate for subsequent molecular dynamics evaluation and future experimental validation.

**Figure 7.**
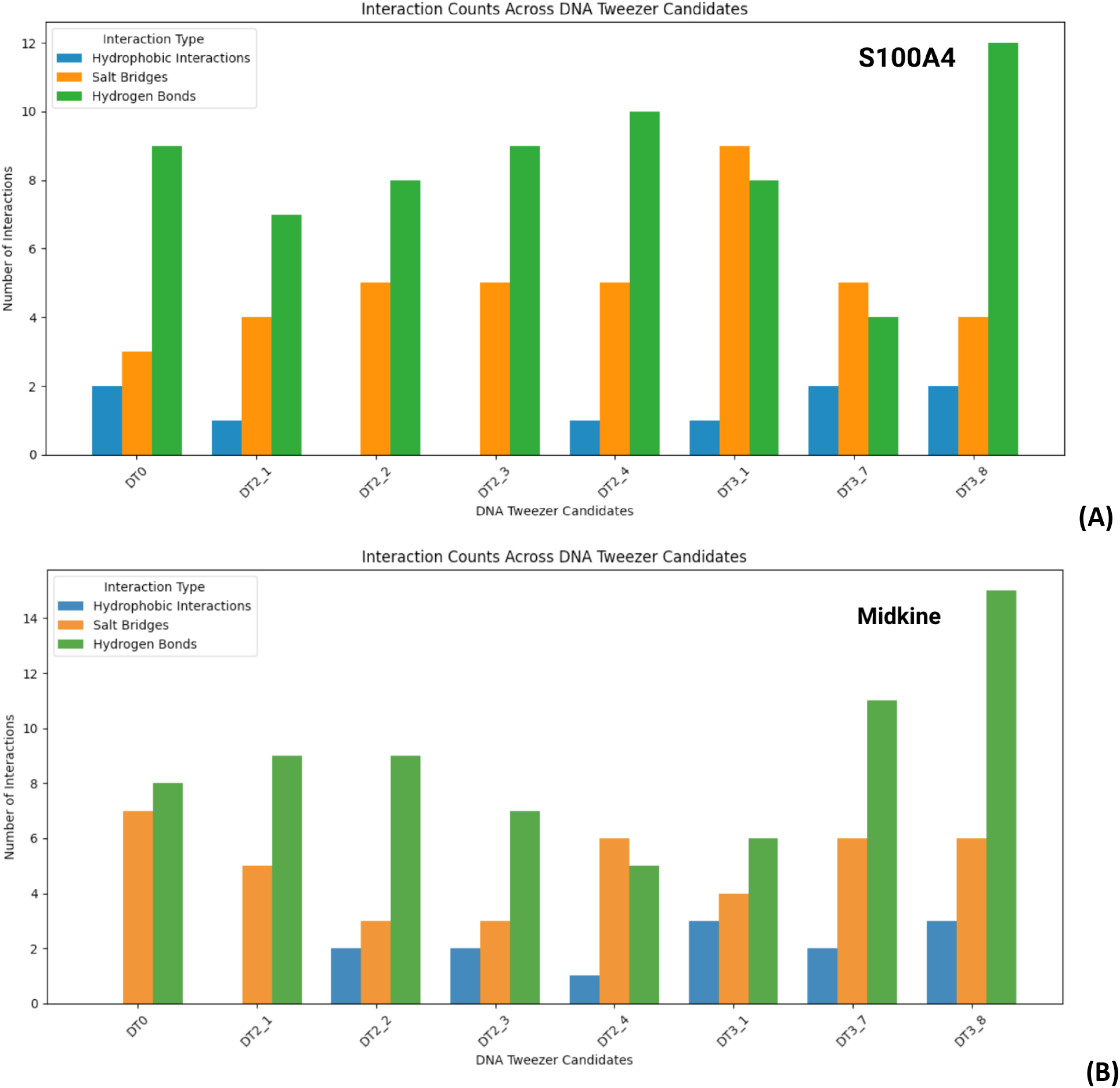
Number of interactions, dealt in major chemical bonds and biochemical interactions formed within the complex between S100A4 and DT3_8 **(A)** and the complex between MDK and DT3_8 **(B)**, with compiled PLIP data. Quantities determined were hydrophobic interactions, salt bridges, and hydrogen bonds. Visualized with Python.

Functionally, DT3_8’s establishment of multiple hydrogen bonds between nucleotide base functional groups and polar residues on the proteins’ surface stabilizes the orientation of the tweezer arms around both S100A4 and MDK, thereby reducing rotational freedom at binding faces. The distribution of hydrogen bonds was concentrated in solvent-accessible regions that allowed complementarily favorable donor-acceptor structure accessibility.

In addition to hydrogen bonding, the hydrophobic contacts predicted between exposed nucleobase rings and nonpolar surface patches of both biomarkers contributed greatly to increased surface complementarity and partial solvent exclusion in the DT3_8 complex. Although DNA is predominantly hydrophilic due to its phosphate backbone, the aromatic bases can participate in van der Waals interactions when appropriately positioned ***[25]***.

Most notably, multiple salt bridges were predicted between negatively charged phosphate groups of the DNA backbone and positively charged lysine and arginine regions on the surface of S100A4 and midkine. These electrostatic interactions are consistent with the central design hypothesis that charge complementarity between DNA strands and a lysine/arginine-enriched protein surface would promote selective association, driving the initial binding event that would be sustained by stronger biochemical interactions later on. Compared with other candidate constructs, DT3_8 exhibited the highest number of predicted salt bridges, clustered around the positively charged C-terminal region of S100A4 and MDK.

### 3.4 Molecular Dynamics and Stability in a Physiological Environment

To assess whether the predicted interactions would remain stable under physiologically relevant conditions—namely, the 3D human body environment with 70% water and concentrations of sodium chloride ions—molecular dynamics simulations were performed using GROMACS (*Figure 8A, 8B)*. RMSF analysis revealed that the regions of the tweezer directly involved in binding exhibited very low fluctuation, suggesting that the binding interface remained rigid and well-maintained over time. The two complexes, DT3_8 / MDK and DT3_8 / S100A4, maintained relatively low RMSD values throughout the simulation, indicating that the complex that both forms has minimal structural drift and strong global stability.

**Figure 8.**
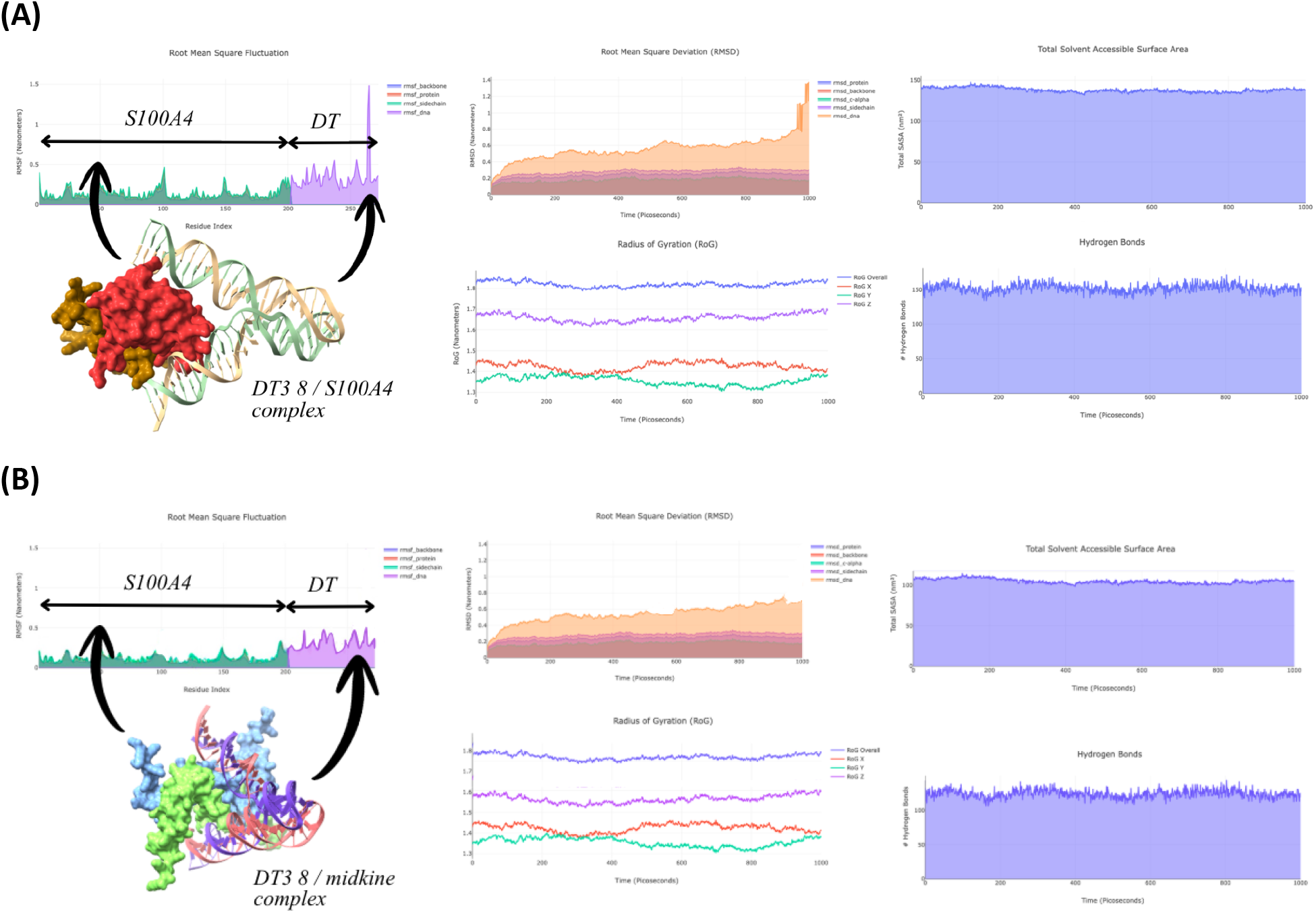
Results of statistical analysis on molecular docking outcomes and viable binding interactions between models of DT3_8 / S100A4 **(A)** and DT3_8 / MDK **(B)** complexes. Both complexes undergo little to no structural fluctuation and retain the surface area necessary for maintaining the integrity of hydrogen bonds.

Notably, the hydrogen bonds identified by PLIP, paired with solvent-accessible surface area, persisted constantly throughout the simulation, demonstrating that the interactions remained stable in a dynamic, water-based environment. This consistency is essential for any diagnostic or theranostic application, as nanodevices must maintain structural integrity *in vivo*, a volatile environment with some extraneous variables.

### 3.5 Overall Interpretation

Across all analyses—structural modeling, docking, interaction profiling, and molecular dynamics—DT3_8 consistently emerged as the strongest candidate in this computational framework. Its superior binding affinity, reproducible docking geometry, high density of stabilizing interactions, and robust dynamic stability collectively support its designation as the leading DNA tweezer for targeting known biomarker proteins S100A4 and midkine.

These results validate the central hypothesis that electrostatic complementarity between negatively charged DNA nanostructures and positively charged GBM biomarkers can be leveraged to design selective, stable, and potentially theranostic nanodevices.

The leading tweezer sequence (DT3_8) was: GGGTATACGTAGGGCTAGGCCAATTTGGCCTAGGCCTAACATACTAG 5’ → 3’

CCCATATGCATCCCGATCCGGTTAAACCGGATCCGGATTGTATGATC 3’ → 5’

## 4. Discussion

The results of this study indicate that DNA tweezers can be computationally designed to potentially recognize and bind the glioblastoma biomarker proteins S100A4 and midkine through a combination of structural complementarity, electrostatic attraction, and stable intermolecular interactions. The strong negative charge of the DNA phosphate backbone aligns naturally with the positively charged lysine- and arginine-rich surfaces of both S100A4 and MDK, providing the initial biochemical foundation for the selective binding event. This principle was confirmed and extended computationally: the DNA tweezer candidate DT3_8 consistently exhibited the most favorable docking energies, the most stable binding orientations, and the highest density of stabilizing interactions across all analyses.

More broadly, this work suggests that DNA tweezers offer a low-cost, programmable, and modular platform for advancing cancer nanotechnology, with particular promise for improving early detection of aggressive tumors such as GBM where infiltration is aggressive and current detection tools fall short. The computational pipeline developed here can be readily adapted to other cancer biomarkers by targeting the same type of protein and adjusting data collection as necessary, highlighting its potential as a scalable framework for future diagnostic and therapeutic development.

While DT3_8 demonstrated strong binding affinity and stability in silico, the translational relevance of these interactions depends on factors such as biomarker concentration, competition with endogenous molecules, and physiological conditions not fully captured in simulation. Compared to existing detection strategies such as antibody-based assays and aptamer systems, DNA nanodevices offer programmability and modularity, though experimental validation is required to assess sensitivity, specificity, and signal generation in biological environments.

### 4.1 Novelty

While DNA nanodevices have been explored for sensing small molecules, ions, and nucleic acids ***[26]***, there are currently no published studies that design DNA tweezers to detect S100A4, MDK, or any other glioblastoma-associated protein. Existing GBM research focuses largely on imaging-based diagnostics, radiomics, or nanoparticle-based contrast agents ***[26]***, none of which operate at the molecular scale to specifically recognize a stimulus. Likewise, although S100A4 and midkine are well-established as markers of GBM invasiveness ***[5, 6, 8]***, no work so far has combined this biomarker with DNA nanotechnology to create an early-stage molecular sensor.

This study fills this gap by computationally designing and evaluating DNA tweezers that can selectively bind either S100A4 or midkine in independent binding events, with a clear framework integrating structure prediction, molecular-level docking, and molecular dynamics simulations into a single study to rationally identify a leading candidate for further investigation in glioblastoma biomarker recognition. This approach is novel because it brings together DNA nanotechnology, two clinically relevant GBM biomarkers, and a systematic, dual-test-based computational design strategy that has not been applied in this context before.

#### 4.1.1 Selectivity and Cross-Reactivity Analysis

To evaluate whether DT3_8 exhibited specificity for S100A4 and MDK rather than generic electrostatic affinity for positively charged proteins, a cross-reactivity panel was constructed using five structurally related, similarly-charge-enriched proteins: S100A1, S100A6, S100B, and calmodulin. Each protein structure was retrieved from UniProt and modeled using AlphaFold under identical parameters as S100A4.

Docking simulations revealed that DT3_8 displayed substantially more favorable ΔG values for S100A4 and MDK than for any other protein in the panel. While all proteins exhibited some degree of electrostatic attraction due to their basic residues, only S100A4 produced a reproducible, low-RMSD binding orientation with a dense network of salt bridges and hydrogen bonds, as suggested by the strongest binding affinity retrieved from HDOCK. In contrast, docking to S100A1, S100A6, S100B, and calmodulin resulted in inconsistent orientations and significantly fewer stabilizing interactions, as indicated by comparably reduced docking scores (*Table 3*).

**Table 3.** Comparative analysis of the pairing between DNA Tweezer candidate DT3_8 and other positively-charged proteins in the same family as S100A4. These complexes show favorable binding but to an extent much less stable and sustained than S100A4, therefore indicating that the surface topology and biochemical interactions add specificity to the binding event between DT3_8 and S100A4 and the binding event of DT3_8 with MDK. Retrieved from HDOCK.

| DNA Tweezers/Protein Complex | Best Docking Score ($\Delta G$) | Best Confidence Score |
| --- | --- | --- |
| DT3_8 / S100A1 | -184.43 | 0.6657 |
| DT3_8 / S100B | -188.41 | 0.6847 |
| DT3_8 / S100A6 | -209.73 | 0.7676 |
| DT3_8 / Calmodulin | -217.68 | 0.7947 |
| DT3_8 / S100A4 | -256.57 | 0.8939 |
| DT3_8 / MDK | -270.84 | 0.9181 |

### 4.2 Challenges

This study was entirely computational, and experimental validation of predicted binding interactions has not yet been performed. Further, the DNA tweezer constructs’ nucleotide sequences were determined and evaluated *in silico*. Their physiological stability, structural integrity, and binding efficiency still require experimental confirmation. For instance, molecular dynamics conducted in this study was only an approximate simulation of the human body environment. It is immensely difficult to account for all extraneous variables and their unpredictable effects on the structure and stability of the DT3_8-protein complex.

### 4.3 Future Directions

Several avenues of future research could build on the findings of this study and advance the DT3_8 construct toward real-world application. First, wet-lab synthesis and fluorescence-based detection assays would be essential for validating the computational predictions ***[27]***. Experimental testing would confirm whether the binding-induced conformational change observed *in silico* produces a measurable fluorescent signal. Second, because GBM is protected by the blood-brain barrier (BBB) ***[2]***, optimization for BBB penetration—potentially through chemical modification, aptamer attachment, or carrier-mediated transport—would be necessary for clinical translation ***[28]***. Third, integration with nanoparticle carriers such as liposomes, gold nanoparticles, or polymeric nanocarriers could enhance stability, circulation time, and targeted delivery in biological systems ***[29]***, in order to evaluate the postulated potential of DNA tweezers to have therapeutic ability in GBM. Finally, the dynamic adaptability of DNA nanotechnology would enable expansion to multi-biomarker detection systems ***[30]***, allowing future devices to sense multiple GBM-associated proteins simultaneously. Such multiplexed platforms could improve diagnostic accuracy, monitor tumor progression more comprehensively, and support personalized treatment strategies. Finally, the structure-guided framework developed in this study could be generalized to other cancer biomarkers beyond S100A4 and midkine, enabling scalable design of programmable DNA nanodevices for broader oncologic applications.

## Notes

### Competing Interest Statement

The authors have declared no competing interest.

